# Histone variant H2BE controls activity-dependent gene expression and homeostatic scaling

**DOI:** 10.1101/2024.11.01.620920

**Authors:** Emily R. Feierman, Alekh Paranjapye, Steven Su, Qi Qiu, Hao Wu, Erica Korb

## Abstract

A cell’s ability to respond and adapt to environmental stimuli relies in part on transcriptional programs controlled by histone proteins. Histones affect transcription through numerous mechanisms including through replacement with variant forms that carry out specific functions. We recently identified the first widely expressed H2B histone variant, H2BE and found that it promotes transcription and is critical for neuronal function and long-term memory. However, how H2BE is regulated by extracellular stimuli and whether it controls activity-dependent transcription and cellular plasticity remain unknown. We used CUT&Tag and RNA-sequencing of primary neurons, single-nucleus sequencing of cortical tissue, and multielectrode array recordings to interrogate the expression of H2BE in response to stimuli and the role of H2BE in activity-dependent gene expression and plasticity. We find that unlike Further, we show that neurons lacking H2BE are unable to mount proper long-term activity-dependent transcriptional responses both in cultured neurons and in animal models. Lastly, we demonstrate that H2BE knockout neurons fail to undergo the electrophysiological changes associated with homeostatic plasticity in neurons after long-term stimulation. In summary, these data demonstrate that H2BE expression is inversely correlated to activity and necessary for long-term activity-dependent responses, revealing the first instance of a histone variant involved in the homeostatic plasticity response in neurons.

## INTRODUCTION

Experience-dependent plasticity is core to brain function, underlying a multitude of processes in the brain, including synapse development, learning and memory, and long-term adaptation to environmental inputs. Neurons have the remarkable capability to change their structural and functional properties in response to diverse environmental stimuli. At a molecular level, transcriptional programs that alter neuronal function and circuitry are critical to multiple forms of plasticity.

Synaptic plasticity can be broadly categorized into two categories: Hebbian plasticity and homeostatic plasticity^1–4^. Hebbian plasticity occurs on the order of seconds to minutes following synaptic activity and is a positive-feedback mechanism leading to strengthening of a given synapse^5,6^. Homeostatic plasticity, which occurs over hours or days, is a negative-feedback process that allows neurons to constrain synaptic signaling to a healthy dynamic range, avoiding runaway potentiation or depression following long-term changes in excitation^7,8^. Hebbian and homeostatic plasticity both require transcription but have distinct activity-dependent transcriptional signatures^1^. Specifically, Hebbian plasticity is marked by the induction of immediate-early genes (IEGs), which are induced within minutes of neuronal activation^9–11^. In contrast, homeostatic plasticity modulates expression of key synaptic proteins, such as postsynaptic receptors, scaffolding proteins, and cell-adhesion molecules via induction of delayed response genes involved in long-term synaptic remodeling^1,12^.

Recent work has revealed that histone variants play a key role in neuronal plasticity^13,14^. Expression of variants H2A.Z and H3.3 in the cortex and hippocampus is regulated *in vitro* in response to pharmacological changes in neuronal activity as well as *in vivo* in mice during learning. Importantly, incorporation and eviction of these variants promotes adaptive activity-dependent gene expression. Prior work demonstrated that the histone variant H2BE is regulated by olfactory stimuli in olfactory sensory neurons. In this unique population, neurons have high rates of turnover and can be replaced by newly generated neurons. Here, H2BE expression is inversely correlated to olfactory receptor activation and controls neuronal survival^15^. Our recent work has shown that H2BE is also expressed throughout the brain where it controls chromatin accessibility, gene expression, and long-term memory^16^. However, whether H2BE is regulated by neuronal activity outside of the olfactory system and how it controls activity-dependent transcription and plasticity remain unknown.

Here, we demonstrate that neuronal activity regulates H2BE expression in the mouse cortex and we define the function of H2BE in regulating activity-dependent gene expression and homeostatic plasticity. First, we show that H2BE expression is inversely regulated by activity in primary cortical neurons and in mouse cortical tissue. Further, using Cleavage Under Targets and Tagmentation (CUT&Tag) with sequencing, we find that neuronal activity results in decreased genomic enrichment of H2BE at neuronal promoters. We then analyzed transcriptional changes in H2BE WT and KO cortical neurons following modulation of neuronal activity and show that H2BE is required for long-term activity-dependent gene expression. Next, we use multielectrode array recordings to demonstrate that H2BE mediates neuronal firing properties both under basal conditions and in response to long-term stimuli. Lastly, we use single-nucleus RNA-sequencing paired with a robust seizure paradigm to demonstrate that H2BE effects on activity-dependent gene expression *in vivo*. Together these data demonstrate that H2BE is a critical mediator of neuronal responses to stimuli and provide the first evidence of a widely expressed mammalian H2B variant capable of linking environmental inputs to activity-dependent transcriptional states.

## RESULTS

### H2BE expression is inversely correlated with neuronal activity

We previously found that H2BE is widely expressed in multiple mouse tissues and particularly abundant in the brain. Further we found that H2BE is critical in controlling chromatin accessibility, gene expression, synaptic strength, and long-term memory in mice^16^. Prior work demonstrated that H2BE is repressed by olfactory receptor activation in the main olfactory epithelium^15^, suggesting it may be controlled by extracellular stimuli. However, whether H2BE is regulated by neuronal activity beyond olfactory receptor activation, and whether H2BE plays a role in regulating activity-dependent gene expression and neuronal responses remains unknown.

Given the high levels of H2BE in the cortex and its established role in regulating chromatin and gene expression under basal conditions, we examined H2BE regulation in cortical neurons. To test whether H2BE is modulated by activity, we pharmacologically manipulated neuronal activity in primary neuronal cultures derived from wildtype (WT) E16.5 embryonic cortices and measured H2BE levels. To increase activity, we used brain-derived neurotrophic factor (BDNF), which acts through binding of TrkB receptors to trigger downstream signaling cascades and by altering neuronal excitability directly^17^. Unlike other histone variants which are typically induced by activity^18–21^, we observed a significant reduction in H2BE transcript levels with increasing time after BDNF treatment (Figure 1A). Strikingly, after 48h, expression of the gene encoding H2BE, *H2bc21,* had reduced to nearly half of baseline levels. This finding extended to a reduction in total H2BE protein levels at 48h (Figure 1B). To ensure that this effect was due to changes in neuronal activity and not specific to BDNF, we performed the same experimental paradigm using the GABA_A_ receptor antagonist bicuculline and observed comparable reductions in *H2bc21* levels, although with distinct kinetics (Supplemental Figure 1A). Lastly, we examined the effect of decreasing neuronal activity to determine if this effect was bidirectional. We treated neurons with a combination of tetrodotoxin (TTX), a Na^+^ channel blocker, and the NMDA receptor antagonist D-2-amino-5-phosphonovalerate (D-AP5) to dampen neuronal activity. In response to TTX/D-AP5, we observed a significant increase in *H2bc21* expression (Figure 1A). Further, using a transcriptional inhibitor, we determined that this effect requires new transcription (Supplemental Figure 1B). However, we did not detect an equivalent increase in H2BE protein levels at 48h, suggesting additional mechanisms are involved in regulating translation or stability of H2BE protein or that extended periods of time are required to detect changes in H2BE protein levels (Figure 1B).

**Figure 1.**
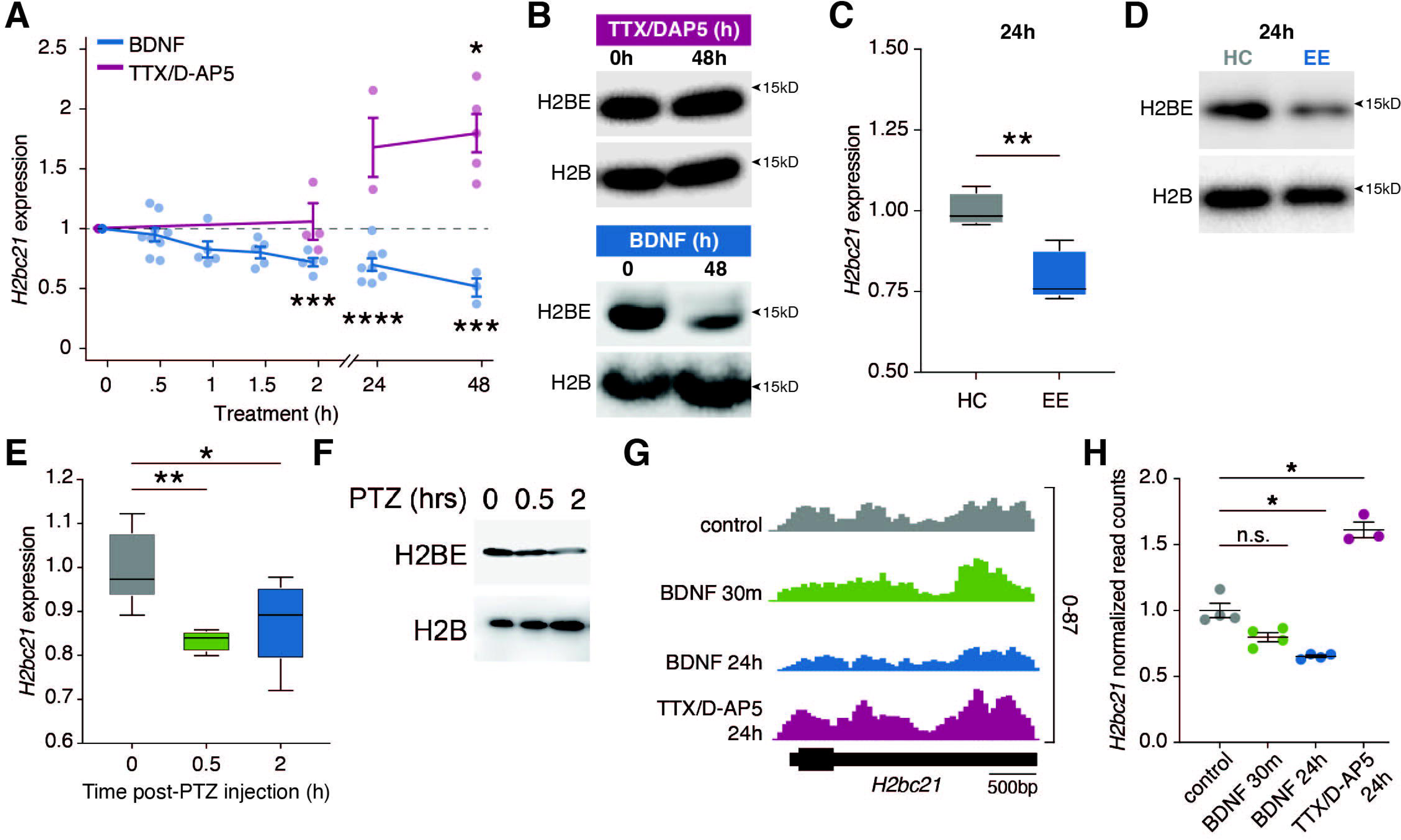
H2BE expression is inversely correlated with neuronal activity. (A) qRT-PCR quantification of H2BE transcript expression following treatment with BDNF or TTX/D-AP5 (n=3-9 biological replicates per timepoint; Kruskal-Wallis test with multiple comparisons). (B) Immunoblot for H2BE and H2B in histone extracts from primary cultured neurons treated with BDNF *(bottom)* or TTX/D-AP5 *(top)*. H2B serves as loading control. (C) qRT-PCR quantification of H2BE transcript expression in mice exposed to an enriched environment (EE) or home cage (HC; n=5 mice per condition; unpaired t-test). (D) Immunoblot for H2BE and H2B in histone extracts from cortical tissue after 24h in EE or HC. H2B serves as loading control. (E) qRT-PCR quantification of H2BE transcript expression in mice following PTZ-induced seizures (n = 4-7 mice per condition; one-way ANOVA with Dunnett’s multiple comparison tests). (F) Immunoblot for H2BE and H2B in histone extracts from cortical tissue after seizure. H2B serves as loading control. (G) RNA-seq gene tracks at *H2bc21*. Scale bar represents 500 base pairs (bp). (H) Normalized read counts within *H2bc21* (n=3-4 biological replicates per condition; one-way ANOVA with Dunnett’s multiple comparison tests). *p<.05, **p<.01, ***p<.001, ****p<.0001, n.s. = not significant.

To assess whether neuronal activity reduces H2BE levels *in vivo* using a physiologically relevant stimulus, we placed mice in an enriched environment (EE) cage for 24 hours. EE is a well-established way to increase neuronal activity, by providing a richer, more stimulating environment than standard housing (reviewed in ^22^). We observed a decrease in H2BE transcript and protein in the cortex of mice exposed to an enriched environment for 24 hours (Figure 1C-D). To further examine the effect of neuronal activation on H2BE levels *in vivo*, we induced seizures using intraperitoneal injection of pentylenetetrazol (PTZ), which causes a rapid and large-scale increase in excitatory activity throughout the brain through inhibition of GABA_A_ receptors. We found that mice injected with PTZ had reduced levels of H2BE transcript and protein by just 2h post-injection compared to control mice that received saline injections, and levels continued to decrease over 8 hours (Figure 1E-F, Supplemental Figure 1C). Together, these data illustrate that H2BE expression is inversely correlated to activity both in primary cultured neurons and *in vivo*.

To further confirm these findings, we performed RNA-sequencing (RNA-seq) on H2BE WT and KO primary cortical neurons following treatment with BDNF for 30m or 24h, as well as TTX/D-AP5 for 24h. We again found that *H2bc21* is significantly downregulated with 24h BDNF treatment and upregulated with TTX/D-AP5 (Figure 1G-H), supporting the finding that H2BE expression is inversely regulated by long-term changes in neuronal activity.

We next sought to determine whether the activity-dependent regulation of H2BE expression affects H2BE enrichment within chromatin or the localization of H2BE throughout the genome. Previous work from our lab showed that H2BE is enriched at promoters of highly expressed synaptic genes^16^. Given this specificity of H2BE localization and the need for neurons to regulate synaptic genes in response to changes in neuronal activity, we hypothesized that activity may affect H2BE enrichment at these sites. To test this, we performed CUT&Tag with sequencing on primary cortical neurons. Because we did not observe a change in H2BE transcript with 30m BDNF or changes in H2B protein levels with 48h TTX/DAP5 treatment, we focused here on the changes in H2BE enrichment at 24h after BDNF treatment. In agreement with our previous work, we found that H2BE is enriched at neuronal promoters (Supplemental Figure 2A-C). Further, we found that H2BE incorporation around transcription start sites is decreased genome-wide with 24h BDNF treatment (Supplemental Figure 2A,C-E). We next specifically examined whether H2BE peaks identified in control conditions change in response to 24h BDNF. We observed a drastic reduction in signal following 24h BDNF within control H2BE peak regions (Figure 2A-B). In fact, 79.6% (1,082) of H2BE peaks were lost following 24h treatment with BDNF with only 311 H2BE peaks gained (Figure 2C). Gene ontology analysis reveals that peaks with decreased H2BE enrichment at promoters following stimulation are related to mRNA processing, autophagy, and key neuronal functions such as intracellular transport and vesicle organization (Figure 2D). Conversely, only two broad terms were significantly enriched within genes with increased H2BE enrichment (Supplemental Figure 2F). We previously found that H2BE enrichment is correlated with gene expression. We therefore examine the relationship between gene expression and H2BE loss following long-term BDNF treatment. Notably, BDNF decreases H2BE at genes regardless of expression level, indicative of a broad response with effects detectable at both high and low expressed genes (Figure 2E-G).

**Figure 2.**
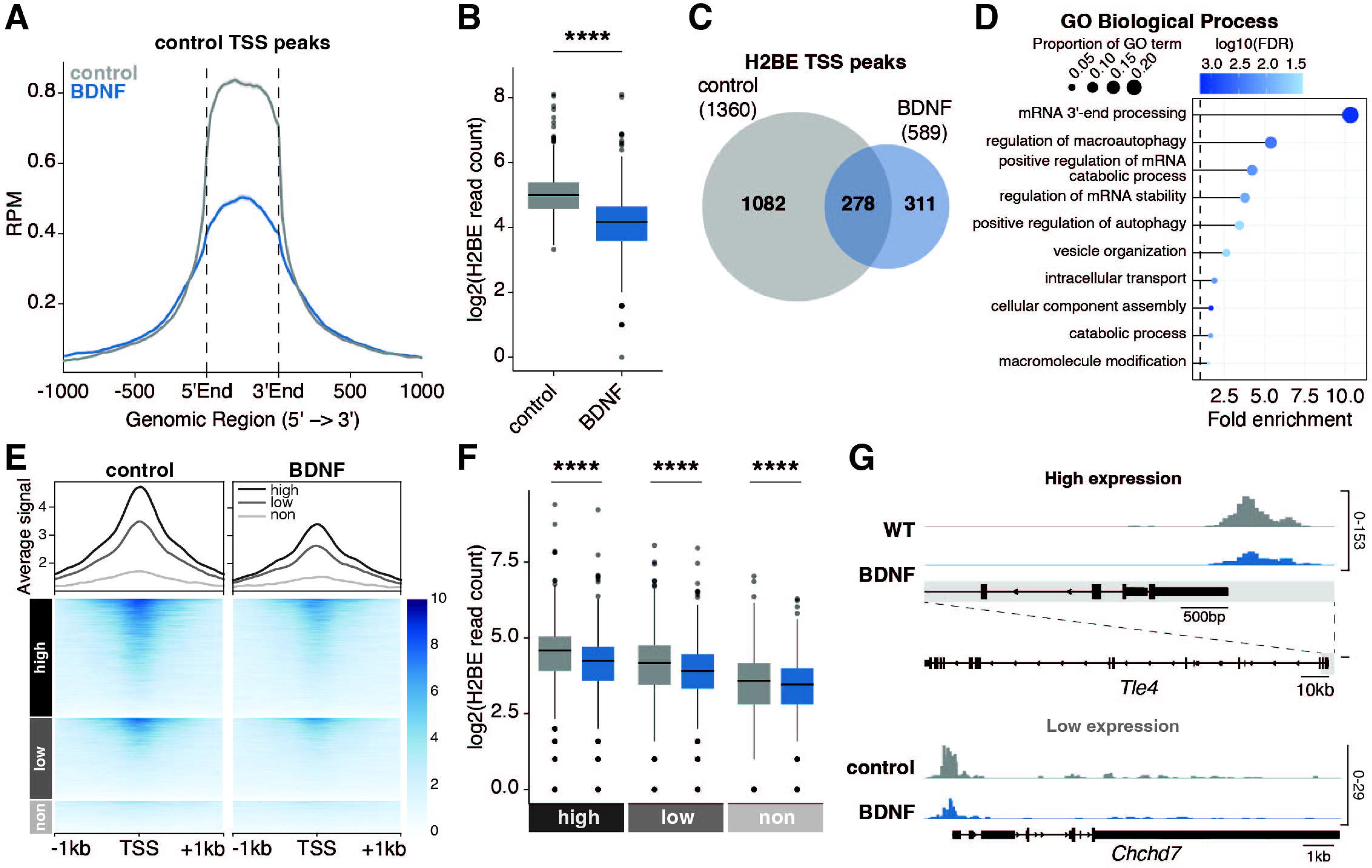
H2BE is downregulated at neuronal promoters following long-term stimulation. (A) Metaplot comparison of CUT&Tag average signal from control and BDNF-treated cortical neurons at all peaks around transcription start sites (TSS) in control. Plot shows read counts per million mapped reads (RPM) at all peaks +/- 1kb (n=3 biological replicates per group). (B) Normalized read counts at all peaks around TSS in control (n = 3 biological replicates per group; unpaired t-test). (C) Overlap of H2BE TSS peak sites in control and BDNF-treated neurons (hypergeometric test; p-adj=2.85x10^-273^). (D) Gene ontology enrichment analysis of H2BE TSS peaks that were downregulated following stimulation (BDNF/control < 0.5). (E) Metaplot comparison *(top),* heat map *(bottom)*, and (F) normalized H2BE read counts of CUT&Tag signal by gene expression. “Not expressed” was defined as genes with mean normalized read counts <3 by RNA-sequencing. Remaining genes were binned into two equally sized groups by mean normalized read counts. (G) CUT&Tag gene tracks at *Chchd7* and *Tle4.* ****p<.0001.

Together, this work demonstrates that H2BE is both downregulated and lost from chromatin following extended periods of increased neuronal activity. These findings indicate an unexpected and inverse relationship between H2BE expression and neuronal stimuli and provides the first evidence of a histone variant that is decreased in response to increased neuronal activity. Notably, the long time scale of this effect is particularly relevant to mechanisms of homeostatic plasticity in which neurons modulate transcriptional output for proper homeostasis following extended periods of prolonged activity.

### H2BE-KO neurons have a dampened transcriptional response to short-term neuronal activation

Our prior work demonstrated that H2BE promotes transcription through an innate ability to promote chromatin accessibility. Given that H2BE is downregulated by neuronal activity, we therefore hypothesized that H2BE will affect the transcriptional response to these same stimuli. To test this, we treated neurons with BDNF and used RNA-seq to capture gene expression changes at two critical timepoints: 30 minutes for short-term primary response activity-dependent gene expression and 24 hours for long-term activity-dependent gene expression.

We previously found that KO neurons have decreased expression of synaptic genes under basal conditions and correspondingly weakened synapses compared to WT neurons^16^. Based on these findings, we hypothesized that KO neurons will show dampened responses to short-term stimuli. Indeed, following 30 minutes of BDNF treatment, 60 genes were upregulated in WT neurons, with a significant enrichment of immediate-early genes, as expected (Figure 3A, Supplemental Figure 3A). In response to the same stimulus, only a subset of these genes (16) was upregulated in KO, and two genes (*Ecrg4, Ptgds*) were modestly downregulated (Figure 3B-C). When directly comparing across genotypes, we saw broad changes in gene expression in KO neurons, with slightly more differentially expressed genes (DEGs) in BDNF-treated neurons compared to basal conditions (1,517 down and 1,372 up following BDNF treatment compared to 1,117 down and 1,047 up under basal conditions) (Supplemental Figure 3B-D). Finally, we used a stringent interaction model to define how genotype and treatment interact to influence gene expression. We identified 22 genes that met these criteria, including multiple immediate-early genes (e.g. *Fosb, Gadd45b, Egr2, Egr4, Dusp5, Nr4a1, Junb, Fos, Jun, Dusp1, Atf3).* These genes had only slightly different levels at baseline (Figure 3D). However, for all 22 genes, there was dramatically reduced activation of these genes in KO compared to WT following 30m treatment with BDNF (Figure 3D-F). These data suggest a blunted activity-dependent response to short-term BDNF treatment. Given that we did not detect changes in H2BE expression at this timepoint (Figure 1A) and give our prior findings of weakened synaptic strength in H2BE KO neurons, this blunted response is most easily explained as a consequence of decreased synaptic response in KO rather than a direct effect of H2BE in mediating short-term activity-dependent gene induction.

**Figure 3.**
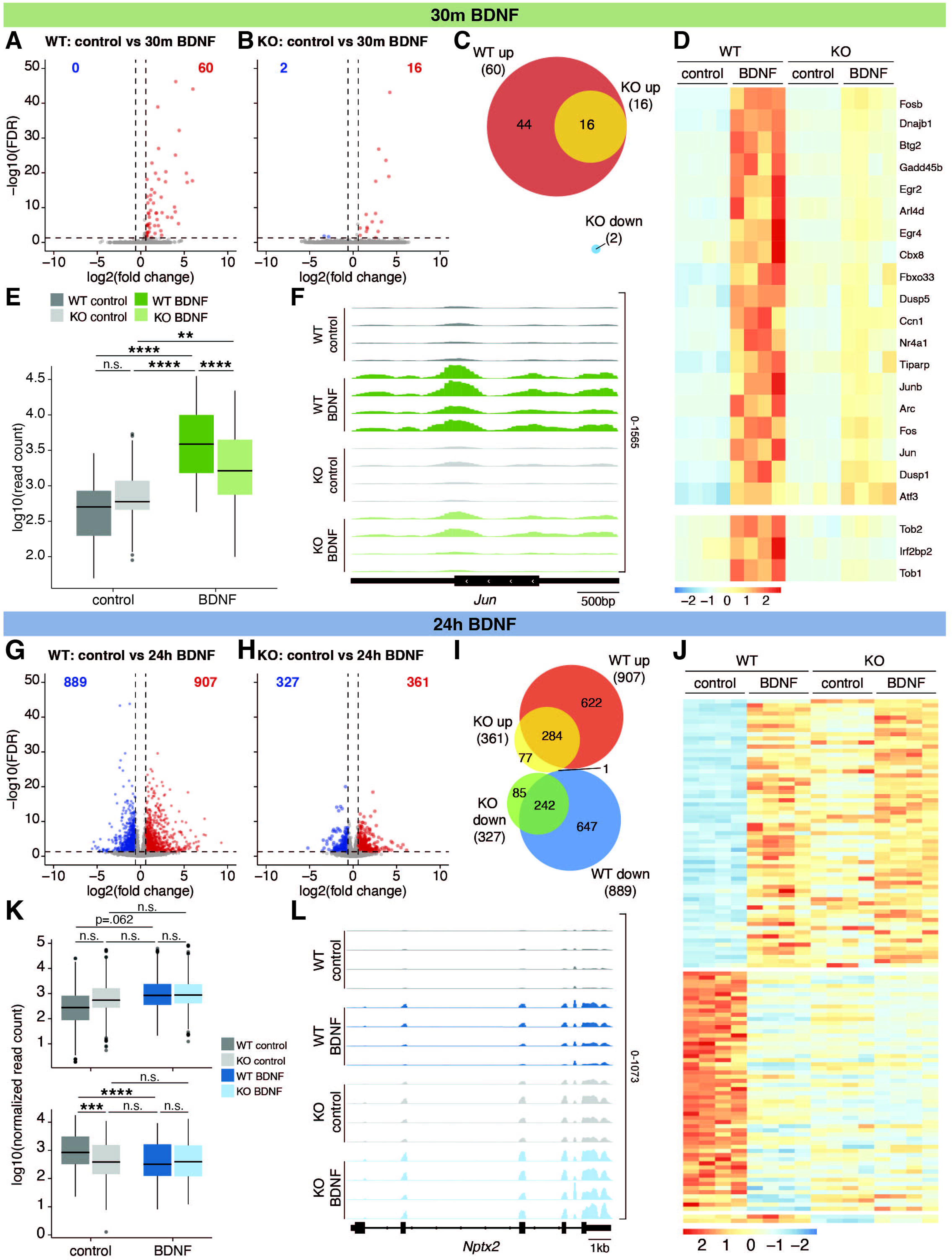
H2BE-KO neurons have a dysregulated transcriptional response to neuronal activation. (A) Volcano plot showing differentially expressed genes (DEGs; FDR<.05, absolute fold change>1.5) between WT and (B) KO cortical neurons +/- 30m BDNF treatment (n=4 biological replicates per group [2 male, 2 female]). Blue=downregulated; red=upregulated. (C) Overlap of DEGs in WT and KO neurons in response to 30m BDNF treatment (hypergeometric test; p-adj=1.69x10^-40^). (D) Heatmap and (E) normalized read counts of genes differentially expressed by an interaction between genotype and treatment (one-way ANOVA and pairwise t-tests with Bonferroni correction). (F) RNA-seq gene tracks for *Jun*. (G) Volcano plot showing DEGs between WT and (H) KO cortical neurons +/- 24h BDNF treatment (n=4 biological replicates per group [2 male, 2 female]). Blue=downregulated; red=upregulated. (I) Overlap of DEGs in WT and KO neurons in response to 24h BDNF treatment (hypergeometric test; down: p-adj=3.18x10^-240^; up: p-adj=3.79x10^-296^. (J) Heatmap and (K) normalized read counts of genes differentially expressed by an interaction between genotype and treatment (one-way ANOVA and pairwise t-tests with Bonferroni correction). (L) RNA-seq gene tracks for *Nptx2*. **p<.01, ***p<.001, ****p<.0001, n.s. = not significant.

### Long-term activity-dependent transcription is dysregulated in H2BE-KO

We next sought to determine whether H2BE plays a role in long-term activity-dependent gene expression. In WT neurons, 24h of treatment with BDNF resulted in 889 downregulated genes and 907 upregulated genes (Figure 3G). KO neurons had a weaker transcriptional response to 24h of BDNF, with 327 downregulated genes and 361 upregulated genes (Figure 3H). There was significant overlap between genes that are differentially expressed in WT and KO neurons, suggesting that KO neurons mount a similar but reduced response to long-term neuronal activation (Figure 3I). Direct comparisons of genotypes under both control and 24h BDNF treatment revealed ∼200 up and down DEGs that were specific to the BDNF condition, while the remainder are in common with control (Supplemental Figure 3B,E-G). Again, using an interaction model, we identified 109 genes that are affected by both treatment and genotype (Figure 3J). Notably, in nearly all cases, expression of these genes at baseline in KO neurons is most similar to WT neurons after 24h BDNF treatment (Figure 3K), including at the scaffold protein *Nptx2* (Figure 3L). These data, along with corresponding activity-dependent decreases in H2BE (Figure 1), support a model in which neurons decrease H2BE to modulate transcription. Thus, KO neurons which are already lacking H2BE appear similar at the transcriptional level to WT neurons following 24h BDNF treatment and fail to undergo further gene expression changes to the same extent as WT neurons.

### H2BE is required for homeostatic plasticity responses following long-term increases in activity

We next utilized a multielectrode array (MEA) recording system to measure activity of WT and KO primary cortical neurons both at baseline and in response to neuronal activation (Figure 4A). We treated neurons with BDNF and measured activity after 48h based on prior literature demonstrating that homeostatic scaling is detectable by MEA recordings at this timepoint^23,24^. After BDNF treatment, WT neurons had significantly decreased firing rate, as well as fewer single-electrode bursts and spikes within bursts (Figure 4B-E). These findings are consistent with prior MEA findings demonstrating similar responses using long-term pharmacological manipulations to increase activity^23,24^. KO neurons showed a similar decrease in spike number to WT (Figure 4D), indicating some activity responses are intact in the absence of H2BE. However, KO neurons also showed multiple divergent responses to long-term BDNF treatment including in several metrics in which they failed to respond (Figure 4B-F). For bursting patterns, control KO neurons appear more similar to stimulated WT neurons, with a significant effect on spikes within bursts and a similar trend in number of bursts (Figure 4E-F). As with transcriptional responses to long-term stimuli, KO neurons appear ‘pre-scaled’ in their bursting patterns and fail to show significant further changes following long-term stimulation. KO neurons also had significantly fewer spikes per burst, and a corresponding increase in inter-spike interval within bursts and overall burst duration (Supplemental Figure 4A-C). Interestingly, while WT neurons downregulate spike amplitude in response to BDNF, KO neurons fail to respond, suggesting that KO neurons are unable to properly scale spike strength in response to prolonged activation even in specific metrics where they are not already at a ‘floor’ (Figure 4F-G). Together, these data show that neuronal activity is dysregulated in KO neurons, both under basal conditions and during synaptic plasticity following long-term neuronal activation. Specifically, we observe 1) metrics for which KO neurons are different under basal conditions, 2) metrics with patterns matching our transcriptomic observations in which control KO neurons appear most similar to WT neurons following a long-term stimulus, and 3) metrics for which only WT neurons, and not KOs, respond to activity. These findings demonstrate that while scaling responses are divergent and likely occur through multiple mechanisms, H2BE is required for specific homeostatic responses to long-term increases in activity.

**Figure 4.**
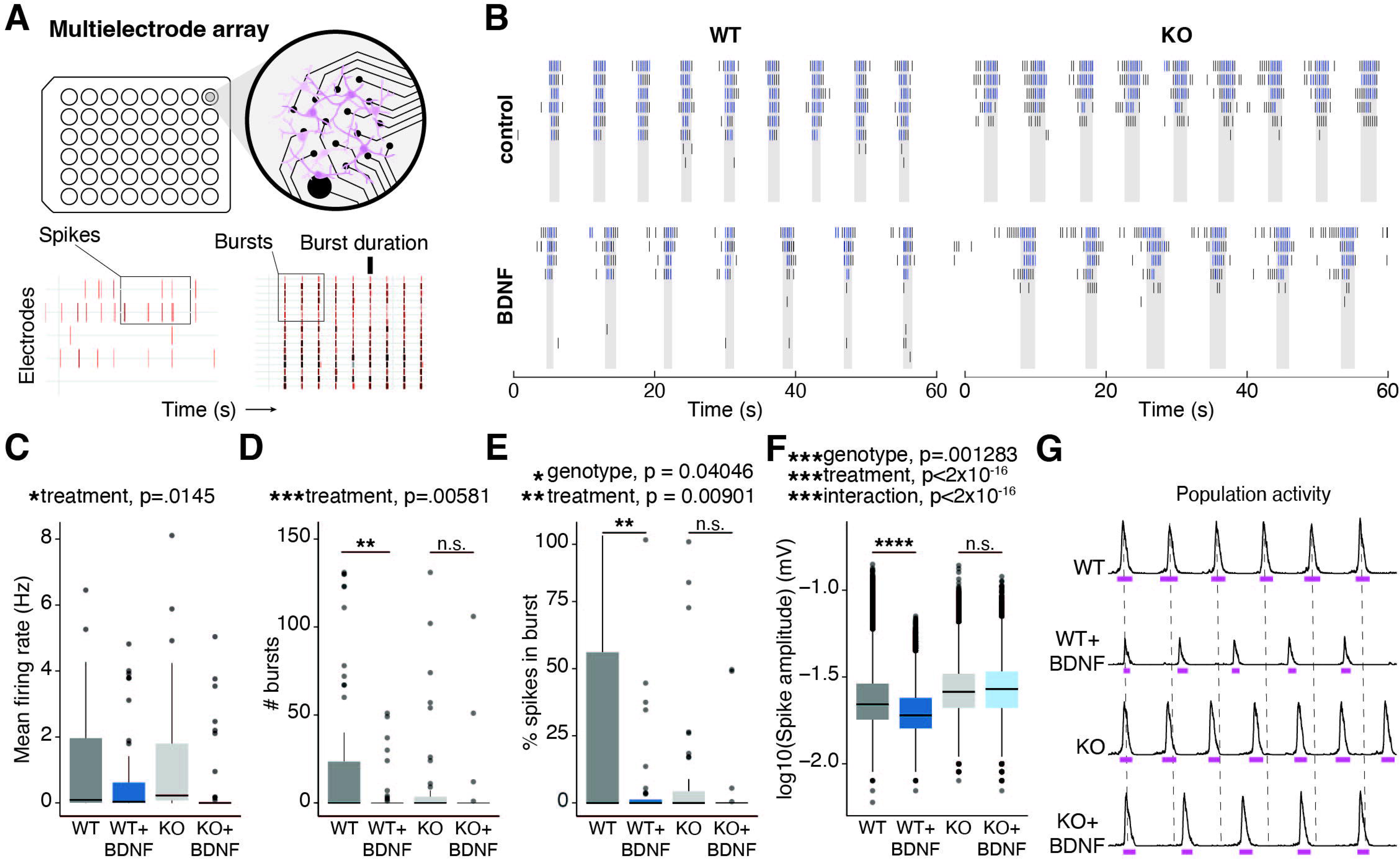
H2BE is required for homeostatic plasticity responses following long-term increases in activity. (A) Schematic of multielectrode array experimental set up. Example raster plots showing spikes and bursts. (B) Firing rate, (C) single-electrode burst count, (D) percent spikes that fell within single-electrode bursts, and (E) mean spike amplitude for WT and KO neurons at 18 days *in vitro* (DIV) (n=3 biological replicates per group, 2 technical replicates per biological replicate). (F) Representative raster plot showing individual spikes and bursting on individual electrodes. (G) Spike count, (H) single-electrode burst count, (I) percent spikes that fell within single-electrode bursts, and (J) mean spike amplitude for WT and KO neurons +/- 48h BDNF at 20 days *in vitro* (DIV) (n=3 biological replicates per group). (K) Representative population activity histograms at 20 DIV. Pink lines represent burst duration. *p<.05, **p<.01, ***p<.001, ****p<.0001, n.s. = not significant.

### Loss of H2BE affects the transcriptomic response to seizures

We next sought to determine whether H2BE plays a similar role in the transcriptional response to activity *in vivo*. We utilized PTZ-induced seizures, an extremely robust method for activity induction, to ensure that even adult KO brains which have reduced synaptic strength^16^ will experience increased neuronal activation. Using a high dose of PTZ (5 mg/mL), we successfully induced seizures in both WT and KO mice. Interestingly, while we did not detect a significant change in seizure severity scores using a Racine scale, fewer KO mice reached higher seizure levels (Supplemental Figure 5A).

To ensure that we analyzed cells that experienced the same degree of seizure, we selected mice that reached a level 4 seizure and performed single-nucleus Drop-seq on cortical tissue. Using Drop-seq, we identified 22 unique cell clusters comprising all expected cell types, namely, 5 inhibitory neuronal clusters, 11 excitatory neuronal clusters, and 6 non-neuronal clusters (oligodendrocytes, oligodendrocyte precursor cells, astrocytes, microglia, radial glial cells, and endothelial cells) (Figure 5A). Fitting with prior findings, there were no major changes in cluster identity across genotypes^16^ although cluster Inh_Rgs9 was under-represented in seizure conditions regardless of genotype (Figure 5A, Supplemental Figure 5B). Differential expression analysis of genotypes at baseline (WTC v KOC) yielded results similar to our previous work, with minor deviations due to the alignment to a modified mm10 genome optimized for 3’ capture in single-cell technology (Supplemental Figure 5C-E). As expected, primary- and secondary-response genes (eg. *Bdnf* and *Nptx2,* respectively) were among the top upregulated genes in seizure conditions for both WT and KO (Figure 5B).

**Figure 5.**
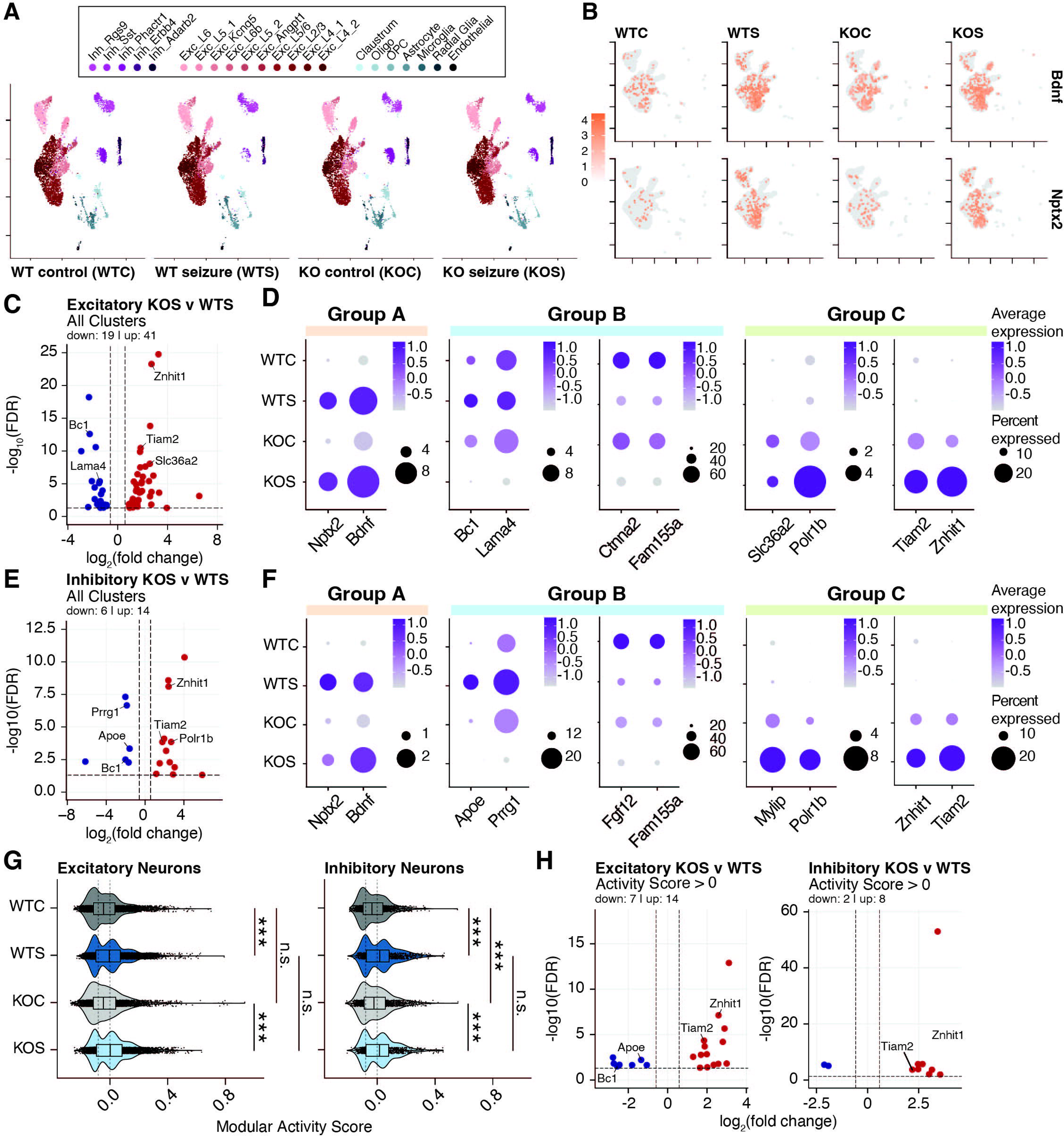
Loss of H2BE affects the transcriptomic response to seizures. (A) UMAP (Uniform Manifold Approximation and Projection for Dimension Reduction) of single-nucleus transcriptomic profiles from adult male (2–4 months) mouse cortices, separated by genotype and seizure status (n=3 biological replicates for WT and KO non-seizure groups, n=2 for WT and KO seizure groups. Each group was randomly subsampled to 7239 nuclei total (nuclei count of smallest group). (B) Feature plots for *Bdnf* and *Nptx2* showing normalized expression between the four experimental groups. (C) Volcano plots of pseudobulk differential gene expression analysis of WTS and KOS in all glutamatergic neuron clusters. (D) Dot plots showing average expression and percent expression of select transcripts in all glutamatergic neurons. (E) Volcano plots of pseudobulk differential gene expression analysis of WTS and KOS in all GABAergic neuron clusters. (F) Dot plots showing average expression and percent expression of select transcripts in all GABAergic neurons. (G) IEG modular activity scores for all glutamatergic or GABAergic neurons by group. (t-tests corrected for multiple testing). Each set was separated into subsets to capture IEG+ and IEG-signatures. (H) Volcano plots for the pseudobulk analysis of KOS versus WTS groups in the IEG+ score subset in excitatory (*left*) or inhibitory (*right*) neurons. ***p<.001, n.s. = not significant.

Direct comparison of genotypes in response to PTZ revealed both down- and upregulated genes in excitatory neurons. When examining all excitatory neuronal clusters, we identified 19 down- and 41 upregulated genes, while individual neuron clusters had more modest changes (Figure 5C, Supplemental Figure 5F). Amongst these changes, we identified three notable patterns of expression regulated by genotype and seizure status: Type A) Common seizure responsive genes with prior elevation in KO; Type B) H2BE-dependent seizure responsive genes (both for up- and downregulated genes; and Type C) genes disrupted by H2BE loss with further aberrant regulation following seizure (Figure 5D). We observed similar but less drastic transcriptional changes when comparing the WT and KO response to PTZ in all inhibitory neurons, with 6 down- and 14 up-regulated genes (Figure 5E, Supplemental Figure 5G). Notably, inhibitory neuron clusters had many of the same DEGs that fell into the three patterns of expression described above, revealing a common set of genes requiring H2BE and/or essential for seizure response across all neuronal subtypes (Figure 5F). Notably, despite the differing stimuli used *in vivo* and in cultured neurons, we similarly detect gene expression responses showing dampened responses in Kos including cases in which control KO neurons most closely resemble WT neurons following a stimulation.

To more precisely parse the role of neuronal activation on transcription *in vivo*, we assigned a Modular Activity Score^25,26^ to all nuclei based on expression of 25 immediate early genes (Figure 5G). We then subset nuclei to specifically compare highly activated neurons (those with a positive activity score) (Figure 5G, Supplemental Figure 5H). We confirmed that this activity score accurately subset nuclei by performing differential expression on nuclei with scores > 0 to those with scores < 0 within genotype for excitatory and inhibitory neurons. All DEGs were upregulated (with one gene significantly downregulated in KOS excitatory neurons), and these genes were almost exclusively immediate early genes, as anticipated (Supplemental Figure 5I-J). We then used this activated population for differential expression analysis to compare activity responses between genotypes following seizure. While these groups contain smaller cell numbers and thus comparisons are less well-powered, we identified 7 down- and 14 upregulated genes in KO excitatory neurons and 2 down- and 8 upregulated genes in KO inhibitory neurons compared to WT (Figure 5H). Notably, this follows similar patterns observed before parsing nuclei based on activity score and thus provides a high-confidence list of genes for which H2BE is necessary to modulate expression following large-scale neuronal activation. Further, this comparison confirms that differential gene expression changes are not due to differences in number of activated neurons by genotype as we detect changes even when specifically examining the population of active neurons. One notable example is *Bc1*, a synaptic non-coding RNA associated with synaptic transmission and dendritic transport^27,28^, which is significantly downregulated in KO neurons following PTZ induction. Amongst upregulated genes, we note *Tiam2*, which has been shown to regulate glutamatergic synaptic transmission and synaptic plasticity^29,30^. Together, these data show that H2BE is necessary for proper regulation of synaptic transmission and plasticity following robust neuronal activation.

## DISCUSSION

Here, we demonstrate the activity-dependent regulation of histone variant H2BE expression and its role in the neuronal response to long-term increases in activity. Unexpectedly, we find that high levels of neuronal activity led to decreased H2BE expression after long-term stimulation, setting H2BE apart from other histone variants in the brain. We show that in neurons lacking H2BE, induction of primary response genes is blunted in response to short-term stimuli. Further, we show that neurons lacking H2BE have dysregulated long-term activity-dependent transcriptional responses both in stimulated primary neuronal cultures and in adult mice with seizures. Lastly, using multielectrode array recordings, we show that KO neurons have aberrant electrophysiological responses to activity that are critical to long-term homeostatic plasticity. Taken together, these data demonstrate that H2BE itself is regulated by neuronal stimuli, and H2BE is required to mount the appropriate transcriptional response to long-term increases in activity.

RNA-seq and MEA data suggest that H2BE plays a role in synaptic scaling in neurons. H2BE levels are decreased with increasing time of stimulation, and knocking out H2BE dampens the transcriptional and electrical changes that are observed in WT neurons following neuronal activation. In fact, both transcriptional and electrophysiological analysis of H2BE-KO suggests that KO neurons are “pre-scaled”, in that at baseline they appear similar to WT levels post-stimulation. The homeostatic scaling response is multi-faceted and requires synaptic remodeling which is ultimately supported by changes in transcriptional state. Recent work demonstrates the importance of chromatin in regulating transcriptional changes involved in synaptic scaling^31,32^. We found that transcriptional plasticity is diminished upon H2BE loss, providing evidence of the first histone variant involved in homeostatic plasticity mechanisms.

While other histone variants, including H2A.Z and H3.3, are also modulated by synaptic activity^13,19,33^, H2BE remains the only variant whose expression is negatively correlated with activity. H2A.Z is actively exchanged in response to neuronal activity; it is evicted from promoters and TSS-flanking CpG islands to allow for the expression of genes and is later reincorporated into chromatin to inactivate those same genes when necessary^18,21,34^. In response to neuronal activation, the variant H3.3 is incorporated at gene bodies and promoters to allow for high expression of active genes^19^. In the case of both H2A.Z and H3.3, these variants accumulate with learning and age. This makes H2BE unique in that its expression increases with age^16^ but decreases with increased activity. Notably, this also suggests that cells such as neurons are capable of using different histone variants to mount proper transcriptional responses to different forms of stimulation.

Notably our findings in cortical neurons are in line with previously identified H2BE regulatory mechanisms in the olfactory system, but result in divergent functional outcomes fitting with the divergent properties of olfactory and cortical neurons, H2BE levels are decreased following activation of olfactory receptors^15^. Olfactory neurons with low activity and high H2BE expression consequently undergo cell death as a means to select for populations of neurons with receptor expression relevant to environmental signals. However, continual cell death and replacement via neurogenesis is a unique feature of the olfactory system that does not occur in the adult cortex. In cortical neurons, we similarly found that H2BE levels are decreased following stimulation (Figure 1). However, we speculated that the functional implications of this activity-dependent regulation are different in the cortex given that there is no neuronal turnover. Instead, we found that H2BE is necessary for the expression of long-term activity-dependent genes and that in the absence of H2BE, neurons fail to undergo appropriate homeostatic scaling responses in response to continued increases in activity. This indicates that multiple tissue types use H2BE to respond to environmental signaling changes but that these responses correspond to different outcomes depending on the functional properties of the cell and tissue.

In summary, this work uncovers the activity-dependent regulation of H2BE in the cortex and the role of H2BE in long-term activity-dependent gene expression (Figure 6). This work provides a novel mechanism by which histone variants in the brain respond to changes in activity to promote homeostatic scaling.

## Acknowledgements

We thank Drs. Catherine Dulac and Stephen Santoro for sharing reagents and mouse lines. E.R.F. was supported by NIH grant F31MH126576. A.P. was supported by NIH grant T32-ES019851. S.S. was supported by UPenn CURF grants.

E.K. was supported by NIH grants 1DP2MH129985, R01NS134755, and R00MH111836 and by the Klingenstein-Simons Fellowship from the Esther A. & Joseph Klingenstein Fund and the Simons Foundation, the Alfred P. Sloan Foundation Research Fellowship (FG-2020-13529), the Brain and Behavior Research Foundation NARSAD Young Investigator Award, and pilot funding from the Epigenetics Institute at the University of Pennsylvania.

## Author Contributions

E.R.F. designed, performed and analyzed the results for most experiments, and wrote the manuscript. A.P. and S.S. performed snDrop-seq analyses. Q.Q. performed snDrop-seq and data preprocessing. H.W. led snDrop-seq experiments. E.K. led the project.

## METHODS

### Mice

An H2BE-KO mouse was generated as described previously^15^. In brief, the endogenous H2BE CDS was replaced with a membrane-targeted mCherry reporter sequence in C57Bl/6 mice. All mice were housed in a 12-hour light-dark cycle and fed a standard diet. All experiments were conducted in accordance with and approval of the IACUC. For all experiments, samples were collected from mice between 3-5-months-old and both male and female mice were included, except for single-nucleus Drop-seq. Drop-seq was performed on cortical tissue from male mice only. This mouse is available from The Jackson Laboratory (strain #023819).

#### Seizure induction

Pentylenetetrazol (PTZ; Sigma P6500) was injected intraperitoneally at 50mg/kg (in PBS). Control mice received PBS injections at equivalent volume. Mice were observed for one hour after injection to score seizures and confirm recovery. The modified Racine scale used to measure seizure induction was as follows:

- Stage 1: Hypoactivity culminating in behavioral arrest with contact between abdomen and the cage.
- Stage 2: Partial clonus (PC) involving the face head or forelimbs.
- Stage 3: generalized clonus (GC) including all four limbs and tail, rearing or falling.
- Stage 4: Generalized Tonic-Clonic seizure (GTC)

Seizure susceptibility score was calculated as: (0.2)(1/PC latency) + (0.3)(1/GC latency) + (0.5)(1/GTC latency). Mice were sacrificed 2 hours after injection.

### Primary neuronal culture

Cortices were dissected from E16.5 C57BL/6J embryos and cultured in supplemented neurobasal medium (Neurobasal [Gibco 21103-049], B27 [Gibco 17504044], GlutaMAX [Gibco 35050-061], Pen-Strep [Gibco 15140-122]) in TC-treated 12- or 6-well plates coated with 0.05 mg/mL Poly-D-lysine (Sigma-Aldrich A-003-E). At 3-4 DIV, neurons were treated with 0.5 µM AraC. For all experiments using cultured cortical neurons, neurons were collected at 12 DIV.

### Pharmacological treatments

The following drugs were diluted into neuronal culture media at the indicated concentrations: brain-derived neurotrophic factor (BDNF; 50 ng/mL, PeproTech 450-02), tetrodotoxin (TTX; 1 µM, Tocris 1069), D-2-amino-5-phosphonovalerate (D-AP5; 100 µM, Tocris 0106), bicuculline (20 µM, Tocris 0130), flavopiridol (FP; 300nM in DMSO, Sigma Aldrich F3055.

### qRT-PCR

cDNA was prepared with a high-capacity cDNA reverse transcription kit (Applied Biosystems 4368813), and quantitative PCR was performed with Power SYBR Green PCR master mix (Applied Biosystems 4367659). Data was analyzed using the common base method^35^.

### Histone extraction

Primary neurons were washed 1X with cold sterile 1X DPBS and collected in 1mL cold 0.4N H_2_SO_4._ Samples were incubated overnight on ice at 4°C. Following the overnight incubation, samples were pelleted for 10 minutes at 18,000g at 4°C and the supernatant was transferred to a new tube. Trichloroacetic acid was added to 25% by volume, and the cells were left on ice at 4°C overnight. Cells were again pelleted 10 minutes at 18,000g at 4°C, and the supernatant was discarded. The pellet was washed 3X with ice-cold acetone. After the third wash, samples were air-dried. The pellet was resuspended in molecular biology-grade H_2_O, incubated at 50°C for 30min, and then sonicated in a Biorupter for 10 min (settings: High, on/off=0.5min/0.5min, 4°C). The 50°C incubation and Biorupter sonication were repeated 1-2X until samples were fully solubilized. Protein concentration was measured using the Bradford assay.

### Western blotting

Histone samples were mixed with 5X Loading Buffer (5% SDS, 0.3M Tris pH 6.8, 1.1mM Bromophenol blue, 37.5% glycerol) and boiled for 10 minutes. Protein was resolved by 16% Tris-glycine SDS-PAGE, followed by transfer to a 0.45-μm PVDF membrane (Sigma-Aldrich IPVH00010) for immunoblotting. Membranes were blocked for 1 hour at RT in 5% milk in 0.1% TBST and probed with primary antibody overnight at 4C. The following primary antibodies were used for western blot analysis: rabbit anti-H2BE (Millipore ABE1384, 1:2000) rabbit anti-H2B (abcam ab1790, 1:10,000). Membranes were incubated with secondary antibody for 1 hour at RT. The following secondary antibody was used for western blotting: goat anti-rabbit HRP (abcam ab6721, 1:5000).

### Environmental enrichment

Wildtype cage-mates were moved into an enriched environment or remained in their standard home cage. Enriched environments consisted of a large rat cage with Alpha-Dri bedding enriched with a tunnel, hut, water dish, Nestlet and other toys to interact with, as well as a vanilla scent. Mice remained in the enriched environment for 24h before tissue was collected.

### CUT&Tag-sequencing

#### Library preparation & sequencing

Input samples were ∼400K primary cortical neurons per biological replicate. CUT&Tag was performed according to published protocols^36^. Concanavalin A-coated beads (Bangs Laboratories BP531) were washed twice with 1mL cold filter-sterilized Binding Buffer (20mM HEPES pH 7.9, 10mM KCl, 1mM CaCl_2_, 1mM MnCl_2_) and resuspended in 11uL/reaction Binding Buffer. Cells were collected in 500uL room temperature Wash Buffer (20mM HEPES pH 7.5, 150mM NaCl, 0.5mM spermidine, supplemented by EDTA-free protease inhibitor [Roche 4693159001]), pelleted at 600 g for 3 min at room temp., washed once with 500uL Wash Buffer (room temp.), pelleted again, and resuspended in 100uL/reaction Wash Buffer (room temp.). To bind cells to ConA beads, 10uL activated beads were added to 100uL cells per reaction, vortexed briefly, and incubated for 10 min at room temperature. Beads were collected on a magnet, supernatant discarded, and beads were resuspended in 50uL ice-cold Antibody Buffer (0.05% digitonin, 2mM EDTA, 0.1% BSA in Wash Buffer). To each reaction, 1ug primary antibody against the target protein was added (anti-H2BE [Millipore ABE1384] or rabbit IgG [Sino Biological CR1], vortexed briefly, and incubated overnight at 4°C on a rocking shaker. Beads were collected on a magnet, supernatant discarded, and beads were resuspended in 50uL secondary antibody (guinea pig anti-rabbit IgG (antibodies-online ABIN101961, diluted 1:100 in Dig-Wash Buffer [0.05% digitonin in Wash Buffer]). Samples were then incubated for 1hr at room temperature on a nutator. On a magnet, beads were washed 2X with 200uL Dig-Wash Buffer. Beads were collected on a magnet, supernatant discarded, and beads were resuspended in 50uL loaded pA-Tn5 adapter complex (Diagenode C01070001-T30, diluted 1:250 in Dig-300 Buffer [20mM HEPES pH 7.5, 300mM NaCl, 0.5mM spermidine, 0.01% digitonin]). Samples were then incubated for 1hr at room temperature on a nutator. On a magnet, beads were washed 2X with 200uL Dig-300 Buffer. Beads were collected on a magnet, supernatant discarded, and beads were resuspended in 300uL Tagmentation Buffer (10mM MgCl_2_ in Dig-300 buffer). Samples were then incubated for 1hr in a 37°C heat block. To stop tagmentation, 10uL 0.5M EDTA, 3uL 10% SDS, and 2.5uL 20mg/mL Proteinase K were added to each reaction and mixed by vortexing. Protein was digested for 1hr in a 55°C heat block. DNA was isolated using Zymo DNA Clean & Concentrator Kit (D4013) and eluted in 22uL molecular biology grade H2O. A universal i5 primer and uniquely barcoded i7 primer were ligated and libraries were amplified by PCR with NEBNext High-Fidelity 2x PCR Master Mix (NEB M0541). Library clean-up was performed with AMPure XP beads (Beckman A63880) and eluted off beads in 25uL Tris-HCl pH 8. Prior to sequencing, library size distribution was confirmed by capillary electrophoresis using an Agilent 4200 TapeStation with high sensitivity D1000 reagents (5067-5585), and libraries were quantified by qPCR using a KAPA Library Quantification Kit (Roche 07960140001). Libraries were sequenced on an Illumina NextSeq550 instrument (42-bp read length, paired end).

#### Data processing and analysis

Reads were mapped to Mus musculus genome build mm10 with Bowtie 2^37^ (v2.4.5). Six million reads from each biological replicate were subset and each condition was then merged across biological replicates (SAMtools^38^ v1.15). Metaplots were generated using ngs.plot^39^ (v2.63). Heatmaps were generated using deepTools^40^ (v3.5.1). Peaks were called using MACS3^41^ (v3.0.0b1) and annotated using Homer^42^ (v4.10). For downstream analysis, we used a Peak Score cutoff of 25 and removed peaks that were assigned to ‘ChrUn’ (unknown chromosome) by Homer. The R package GenomicDistributions^43^ (v1.6.0) was used to analyze the genomic distribution of peaks. IGV tools^44^ (2.12.3) was used to generate genome browser views.

To compare CUT&Tag signal to gene expression, normalized read counts from RNA-sequencing of WT primary neuronal cultures (see RNA-sequencing methods section) were used to generate gene lists by expression level. Genes with base mean < 3 were defined as ‘not expressed’. The remaining genes were divided into 2 bins (by base mean) to define ‘low expression’ and ‘high expression’.

#### Gene ontology

For gene ontology analysis, gene names were assigned to peak coordinates using Homer. PANTHER^63,75^ (v18.0) was used to perform an overrepresentation test against the biological process complete ontology using default parameters. The Mus musculus genome was used as a background gene list. For conciseness and visualization, parent terms were excluded and only the most specific GO terms were plotted.

### RNA-sequencing

#### Library preparation & sequencing

RNA was isolated from primary cortical neurons using Zymo Quick-RNA Miniprep Plus Kit (R1057). Prior to library preparation, RNA integrity was confirmed using an Agilent 4200 TapeStation with high sensitivity RNA reagents (5067-5579). Sequencing libraries were prepared using the TruSeq Stranded mRNA kit (Illumina 20020595). Prior to sequencing, library size distribution was confirmed by capillary electrophoresis using an Agilent 4200 TapeStation with high sensitivity D1000 reagents (5067-5585), and libraries were quantified by qPCR using a KAPA Library Quantification Kit (Roche 07960140001). Libraries were sequenced on an Illumina NextSeq1000 instrument (66-bp read length, paired end).

#### Data processing and analysis

Reads were mapped to *Mus musculus* genome build mm10 with Star^45^ (v2.7.9a). The R packages DESeq2^46^ (v1.38.3) and limma (v3.54.2) via edgeR^47^ (v3.40.2) were used to perform differential gene expression analysis. We defined genes as differentially expressed where FDR<0.05 and absolute fold change >= 1.5. Volcano plots were generated using VolcaNoseR^48^. IGV tools^44^ (2.12.3) was used to generate genome browser views. Overlap significance of gene lists was determined by hypergeometric testing. Differential gene expression analysis for interactions were performed using DESeq2. The input design matrix for model fitting factored genotype and treatment (design=∼genotype + treatment + genotype:treatment). The DESeq function estimates size factors using the standard mean ratio and parametric dispersion fitting. Differential expression was calculated using a generalized linear model with negative binomial GLM fitting accounting for individual effects and the interaction term. Genes exhibiting a significant interaction from the Wald test were extracted from the results table using a contrast specifying the interaction term (0,0,0,-1).

#### Gene Set Enrichment Analysis (GSEA)

The R package FGSEA^49^ was used to perform pre-ranked gene set enrichment analysis (GSEA)^50,51^ based on log2 fold changes obtained from DESeq2 differential expression analysis. The immediate-early gene set was defined in Tullai et al. 2007^52^.

### Multi-electrode array (MEA)

Neurons were plated on CytoView MEA 48-well plates (Axion BioSystems M768-tMEA-48W). Prior to plating, plates were coated with 50 ug/mL poly-D-lysine (Sigma-Aldrich A-003-E) in borate buffer pH 8.4, incubated overnight at 37°C, washed 4X with H_2_O, and air dried overnight. After air drying, wells were coated with 20 ug/mL laminin (Roche 1. 11243217001) in ice-cold Opti-MEM (Gibco 51985091) and incubated 4h at 37°C. Immediately prior to seeding, laminin was removed from wells and 80K neurons were seeded on the CytoView plate. Recordings were performed using an Axion Maestro Pro™ multiwell microelectrode array with 5% CO_2_ at 37°C. Baseline recordings were taken after 18 days *in vitro*. Following baseline recordings, BDNF was added to the wells, and the MEA plate was returned to the incubator for an additional 48h. AxIS software (Axion Biosystems) was used for the extraction of spikes and bursts. Burst activity was defined as a minimum of 5 spikes with a maximum inter-spike interval of 100 ms^53^.

### Single-nucleus Drop-sequencing (snDrop-seq)

#### Nuclei isolation

Snap-frozen brain tissues were homogenized in 1mL Buffer A (0.25M sucrose, 50mM Tris-HCl pH7.4, 25mM KCl, 5mM MgCl2 supplemented by EDTA-free protease inhibitor [Roche 4693159001]) using a pre-chilled dounce and pestle. Homogenate was then transferred to a pre-chilled 15mL conical tube and mixed with 6mL Buffer B (2.3M sucrose, 50mM Tris-HCl pH7.4, 25mM KCl, 5mM MgCl2). An additional 2mL Buffer A was used to rinse leftover homogenate from the dounce and combined with the sample. The homogenate was gently transferred to a pre-chilled 15mL ultracentrifuge tube containing 2mL Buffer C (1.8M sucrose, 50mM Tris-HCl pH7.4, 25mM KCl, 5mM MgCl2). Nuclei were pelleted at 100,000 x g for 1.5hr at 4C using a SWI41 rotor. The supernatant was discarded and 1.5mL Buffer D (0.01% BSA in 1X PBS with 0.5U/uL RNase inhibitor [Lucigen 30281-2]) was gently added to the nuclei pellet and incubated on ice 20min. The nuclei pellet were resuspended and the suspension was transferred to a 1.5mL lo-bind tube.

#### Library preparation and sequencing

The single-nucleus suspensions were individually diluted to a concentration of 100 nuclei/mL in DPBS containing 0.01% BSA. Approximately 1.5 mL of this single-nucleus suspension was loaded for each sNucDrop-seq run. The single-nucleus suspension was then co-encapsulated with barcoded beads (ChemGenes) using an Aquapel-coated PDMS microfluidic device (mFluidix) connected to syringe pumps (KD Scientific) via polyethylene tubing with an inner diameter of 0.38mm (Scientific Commodities)<u>68</u>. Barcoded beads were resuspended in lysis buffer (200 mM Tris-HCl pH8.0, 20 mM EDTA, 6% Ficoll PM-400 (GE Healthcare/Fisher Scientific), 0.2% Sarkosyl (Sigma-Aldrich), and 50 mM DTT (Fermentas; freshly made on the day of run) at a concentration of 120 beads/mL. The flow rates for nuclei and beads were set to 4,000 mL/hr, while QX200 droplet generation oil (Bio-rad) was run at 15,000 mL/hr. A typical run lasts 20 min. Droplet breakage with Perfluoro-1-octanol (Sigma-Aldrich), reverse transcription and exonuclease I treatment were performed, as previously described,<u>29</u> with minor modifications. For up to 120,000 beads, 200 μL of reverse transcription (RT) mix (1x Maxima RT buffer (ThermoFisher), 4% Ficoll PM-400, 1 mM dNTPs (Clontech), 1 U/mL RNase inhibitor, 2.5 mM Template Switch Oligo (TSO: AAGCAGTGGTATCAACGCAGAGTGAATrGrGrG), and 10 U/ mL Maxima H Minus Reverse Transcriptase (ThermoFisher)) were added. The RT reaction was incubated at room temperature for 30min, followed by incubation at 42C for 120 min. To determine an optimal number of PCR cycles for amplification of cDNA, an aliquot of 6,000 beads was amplified by PCR in a volume of 50 μL (25 μL of 2x KAPA HiFi hotstart readymix (KAPA biosystems), 0.4 μL of 100 mM TSO-PCR primer (AAGCAGTGGTATCAACGCAGAGT, 24.6 μL of nuclease-free water) with the following thermal cycling parameter (95C for 3 min; 4 cycles of 98C for 20 sec, 65C for 45 sec, 72C for 3 min; 9 cycles of 98C for 20 sec, 67C for 45 sec, 72C for 3 min; 72C for 5 min, hold at 4C). After two rounds of purification with 0.6x SPRISelect beads (Beckman Coulter), amplified cDNA was eluted with 10 μL of water. 10% of amplified cDNA was used to perform real-time PCR analysis (1 μL of purified cDNA, 0.2 μL of 25 mM TSO-PCR primer, 5 μL of 2x KAPA FAST qPCR readymix, and 3.8 μL of water) to determine the additional number of PCR cycles needed for optimal cDNA amplification (Applied Biosystems QuantStudio 7 Flex). We then prepared PCR reactions per total number of barcoded beads collected for each sNucDrop-seq run, using 6,000 beads per 50-μL PCR reaction, and ran the aforementioned program to amplify the cDNA for 4 + 10 to 12 cycles. We then tagmented cDNA using the Nextera XT DNA sample preparation kit (Illumina, FC-131-1096), starting with 550 pg of cDNA pooled in equal amounts, from all PCR reactions for a given run. Following cDNA tagmentation, we further amplified the tagmented cDNA libraries with 12 enrichment PCR cycles using the Illumina Nextera XT i7 primers along with the P5-TSO hybrid primer (AATGATACGGCGACCACCGAGATCTACACGCCTGTCCGCGGAAGCAGTGGTATCA ACGCAGAGT∗A∗C).<u>68</u> After quality control analysis by Qubit 3.0 (Invitrogen) and a Bioanalyzer (Agilent), libraries were sequenced on an Illumina NextSeq 500 instrument using the 75-cycle High Output v2 Kit (Illumina). We loaded the library at 2.0 pM and provided Custom Read1 Primer (GCCTGTCCGCGGAAGCAGTGGTATCAACGCAGAGTAC) at 0.3 mM in position 7 of the reagent cartridge. The sequencing configuration was 20 bp (Read1), 8 bp (Index1), and 60 bp (Read2).

#### Data preprocessing

Paired end sequencing reads were processed using 10X Genomics Cellranger v5.0.1. Reads were aligned to the mm10 genome optimized for single cell sequencing through a hybrid intronic read recovery approach (https://doi.org/10.1038/s41592-023-02003-w). In short, reads with valid barcodes were trimmed by TSO sequence, and aligned using STAR v2.7.1 with MAPQ adjustment. Intronic reads were removed and high-confidence mapped reads were filtered for multimapping and UMI correction. Empty GEMs were also removed as part of the pipeline. Initial dimensionality reduction and clustering was performed prior to processing to enable batch correction and removal of cell free mRNA using SoupX (https://doi.org/10.1093/gigascience/giaa151). Raw expression matrices with counted, individual nuclei UMI and genes were used for subsequent steps and filtering by QC metrics.

#### Clustering and merging by condition and comparison

Raw matrices for each individual replicate per condition were converted to Seurat objects using Seurat 5.0.1 and filtered to remove UMIs with thresholds of > 200 minimum features, > 250 genes detected per nuclei, < 20% mitochondrial reads, and < 2% ribosomal reads. Replicates were merged to generate objects per condition for the subsequent steps. Each dataset was normalized using the default scale factor of 10000, variable selection was performed using 2000 features, then scaled and centered using all features without regressing any variables. Dimensionality reduction with PCA used the first 30 principal components and the nearest-neighbor graph construction used the first 10 dimensions. Clustering was next performed using a resolution of 0.4 before layers corresponding to each replicate were integrated using CCAIntegration with a k weight of 60 and then rejoined. The dataset per condition was then dimensionally reduced using the integrated CCA at with 30 dimensions and the same resolution of 0.4. ScType (https://doi.org/10.1038/s41467-022-28803-w) was used for automated, de novo cell type identification of the clusters followed by manual curation for clusters with low confidence scores. Each automatically assigned cluster was manually validated using previously generated cluster identity labels^16^. Differential clustering analysis was performed using the scProportionTest R package. For all comparisons, the objects per condition were merged and processed using the same integration methodology above to scale and normalize between all incorporated samples. Pseudobulk differentially expressed genes were identified using DElegate (https://github.com/cancerbits/DElegate?tab=readme-ov-file), a wrapper for EdgeR on single nuclei data, with a fold change threshold of 1.5 and Benjamini-Hochberg adjusted p value ≤ .05. Counts were aggregated by individual mice per condition under the orig.ident identity, then pairwise comparisons were computed using quasi-likelihood dispersion with glmQLFit through the findDE function.

#### Modular activity scoring and subsetting

Modular activity scores were calculated for excitatory and inhibitory neurons using AddModuleScore with the list of the 25 putative immediate early genes (*Arc*, *Bdnf*, *Cdkn1a*, *Dnajb5*, *Egr1*, *Egr2*, *Egr4*, *Fos*, *Fosb*, *Fosl2*, *Homer1*, *Junb*, *Nefm*, *Npas4*, *Nr4a1*, *Nr4a2*, *Nr4a3*, *Nrn1*, *Ntrk2*, *Rheb*, *Sgsm1*, *Syt4*, *Vgf*) against a control feature score of 5. Nuclei with an activity score over zero were isolated as IEG+. Those under the 25^th^ percentile threshold were marked as IEG-. Each IEG score group was remerged and RNA layers were jointed before FindMarkers was used to perform all IEG+/IEG-pairwise comparisons with findDE.

#### Gene ontology

Gene ontology analysis was performed using gProfiler g:GOSt^54^. Each gene list was analyzed using an over-representation test against the gene ontology biological process database with a Benjamini-Hochberg FDR correction for multiple testing correction. Only terms with a size between 0 to 2000 genes were selected for specificity.

---

**Supplemental Figure 1. H2BE transcript expression in neurons following treatment with bicuculline.** (A) qRT-PCR quantification of H2BE transcript expression following treatment with bicuculline (n=2 biological replicates per timepoint; Kruskal-Wallis test). (B) qRT-PCR quantification of H2BE transcript expression following treatment with TTX/D-AP5, flavopiridol (FP), or both (n=3 biological replicates per timepoint; one-way ANOVA with Tukey’s multiple comparisons test). (C) qRT-PCR quantification of H2BE transcript expression in mice exposed to an enriched environment (EE) or home cage (HC) for 1 hour per day for 30 consecutive days (n=4 HC, 5 EE mice; unpaired t-test). (E) Immunoblot for H2BE and H2B in histone extracts from cortical tissue after seizure. H2B serves as loading control. **p<.01, ***p<.001, n.s. = not significant.

**Supplemental Figure 2. CUT&Tag analysis of H2BE in response to neuronal activation.** (A) Metaplot comparison of CUT&Tag average signal from control and BDNF-treated cortical neurons. Plot shows read counts per million mapped reads (RPM) between the transcription start site (TSS) and transcription end site (TES) +/- 2kb (n=3 biological replicates per condition). (B) Genomic distribution of control H2BE enrichment sites relative to the mouse genome (Chi-square test). (C) Distribution of H2BE enrichment sites relative to the nearest TSS. (D) Metaplot comparison of CUT&Tag average signal and (E) normalized H2BE read counts at all transcription start sites. Plot shows read counts per million mapped reads (RPM) around the transcription start site (TSS) +/- 2kb (n = 3 biological replicates per condition). (F) Gene ontology enrichment analysis of TSS peaks gained in the BDNF condition. ***p<.001, ****p<.0001; n.s. = not significant.

**Supplemental Figure 3. Analysis of H2BE effect on the transcriptional response to neuronal activation.** (A) GSEA analysis of genes upregulated in WT following 30m of BDNF treatment. The 60 upregulated genes in WT neurons following 30m of BDNF treatment compared with immediate-early genes, as defined in Tullai et al. 2007. (B) Volcano plot of WT and KO cortical neurons at baseline and (C) after 30m BDNF treatment. (D) Overlap of DEGs in WT and KO neurons at baseline and after 30m BDNF (hypergeometric test; down: p-adj=0; up: p-adj=0). (E) Volcano plot of WT and KO cortical neurons after 24h BDNF treatment. (F) Overlap of DEGs in WT and KO neurons at baseline and after 24h BDNF (hypergeometric test; down: p-adj=0; up: p-adj=0). (G) Gene ontology enrichment analysis of DEGs in KO after 24h BDNF.

**Supplemental Figure 4. MEA analysis of H2BE effect on the neuronal response to BDNF.** (A) Burst duration of WT and KO neurons at 18 days *in vitro* (DIV) (n=3 biological replicates per group, 2 technical replicates per biological replicate). (B) Mean number of spikes per burst for WT and KO neurons +/- 48h BDNF at 20 days *in vitro* (DIV) (n=3 biological replicates per group). (C) Mean inter-spike interval (ISI) for WT and KO neurons +/- 48h BDNF at 20 DIV (n=3 biological replicates per group). (D) Burst duration of WT and KO neurons +/- 48h BDNF at 20 DIV (n=3 biological replicates per group. *p<.05, ***p<.001, n.s. = not significant.

**Supplemental Figure 5. Validation of H2BE KO differential gene expression.** (A) Percentage of WT and KO mice who reached Racine levels 1-4 following PTZ-injection. (B) Analysis of the proportion of control and BDNF nuclei in each cluster. Data represents the fold change in WT vs KO nuclei in each cluster and a confidence interval for the magnitude difference. Clusters with fold difference >1 and FDR<.05 highlighted in pink. (C) Number of differentially expressed genes within each cluster for all excitatory and inhibitory neuronal clusters using the 3’ capture optimized mm10 genome. (D) Volcano plot and gene ontology enrichment analysis of downregulated genes in cluster Ex_L2/3. (E) Volcano plot and gene ontology enrichment analysis of downregulated genes in cluster Inh_Phactr1. (F) Volcano plots of pseudobulk differential gene expression analysis of WTS and KOS in clusters corresponding to cortical layer 2/3, Kcnq5+ neurons, or cortical layer 6. (G) Volcano plots of pseudobulk differential gene expression analysis of WTS and KOS in Sst+ neurons, or Phactr1+ neurons. (H) IEG modular activity scores for all glutamatergic or GABAergic neurons by group. (t-tests corrected for multiple testing). Each set was separated into subsets to capture IEG+ and IEG- signatures. (I-J) Volcano plots for the pseudobulk analysis of either WTS or KOS, comparing nuclei with high modular activity score versus low scores in excitatory (I) or inhibitory (J) neurons.

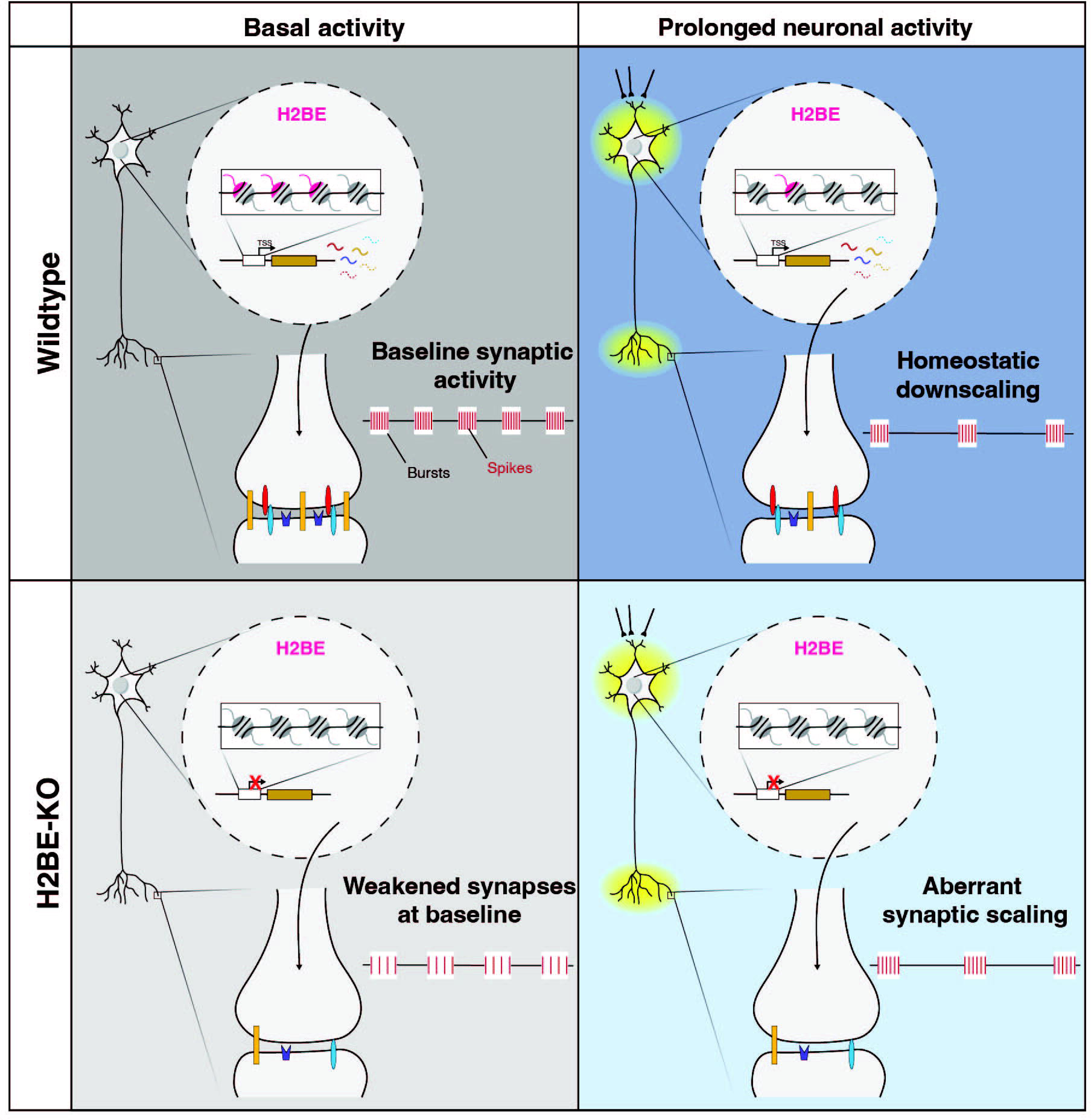

