## Supplementary figures and images for "Histone variant H2BE controls activity-dependent gene expression and homeostatic scaling"

### Supplemental Figure 1

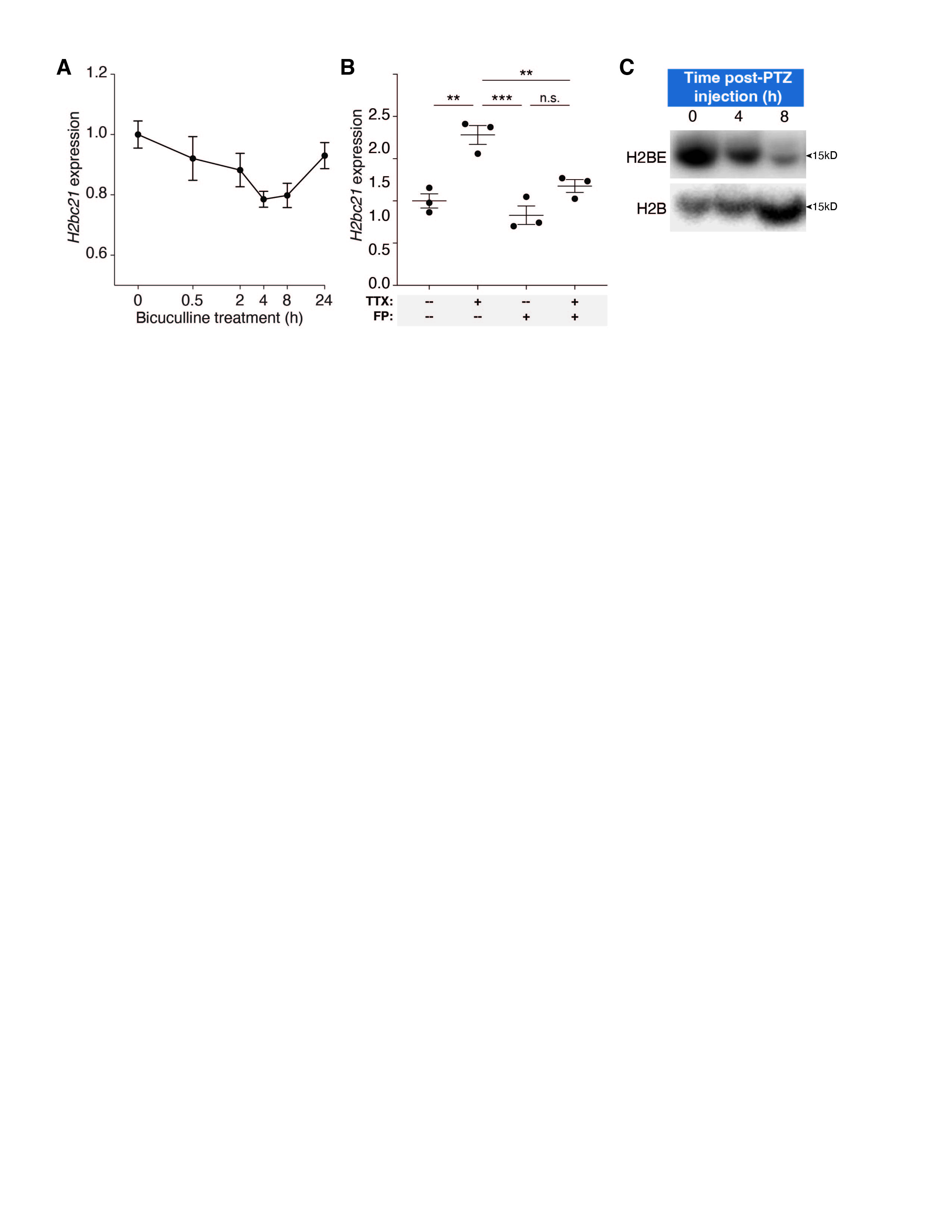

### Supplemental Figure 2

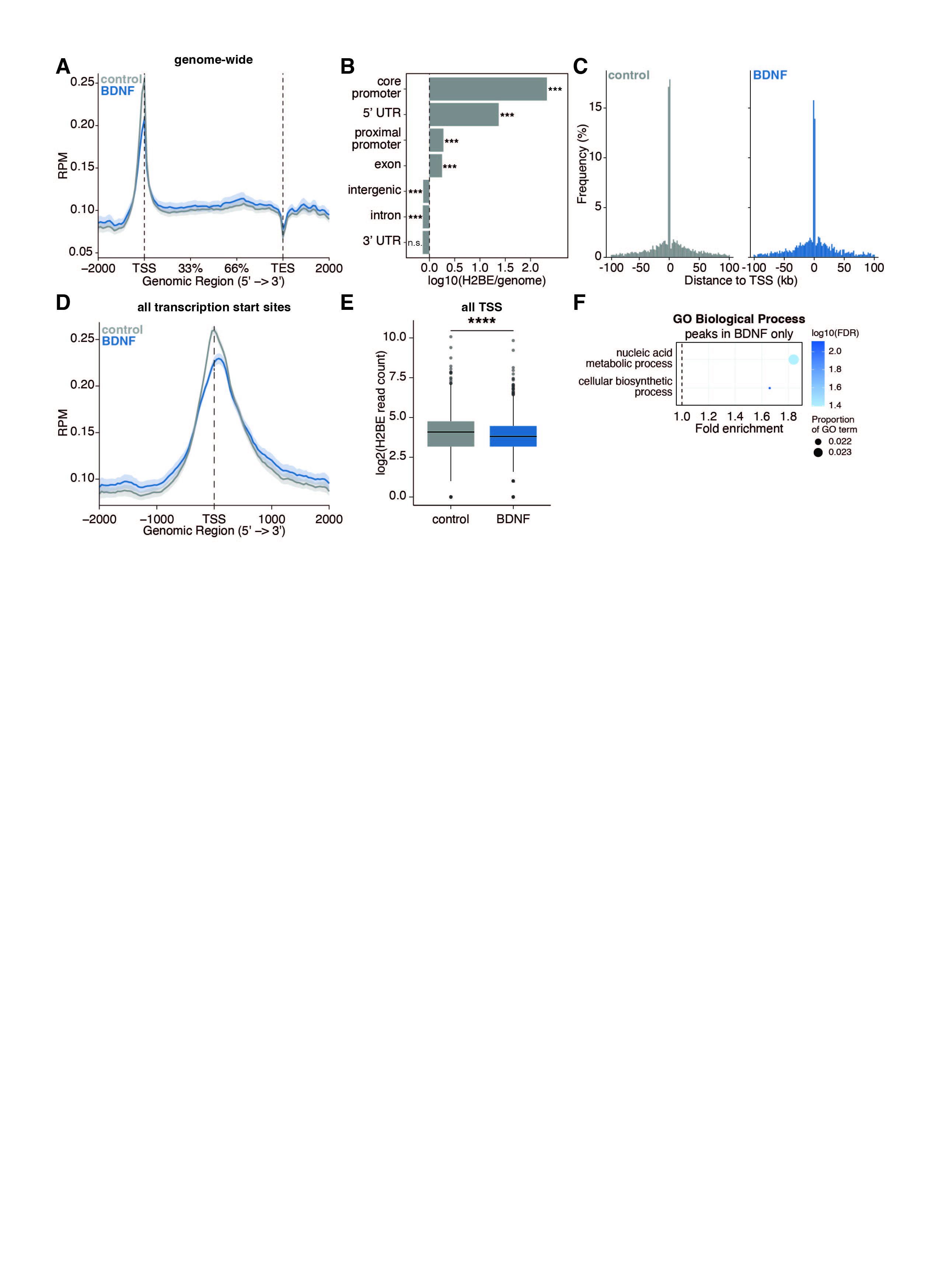

### Supplemental Figure 3

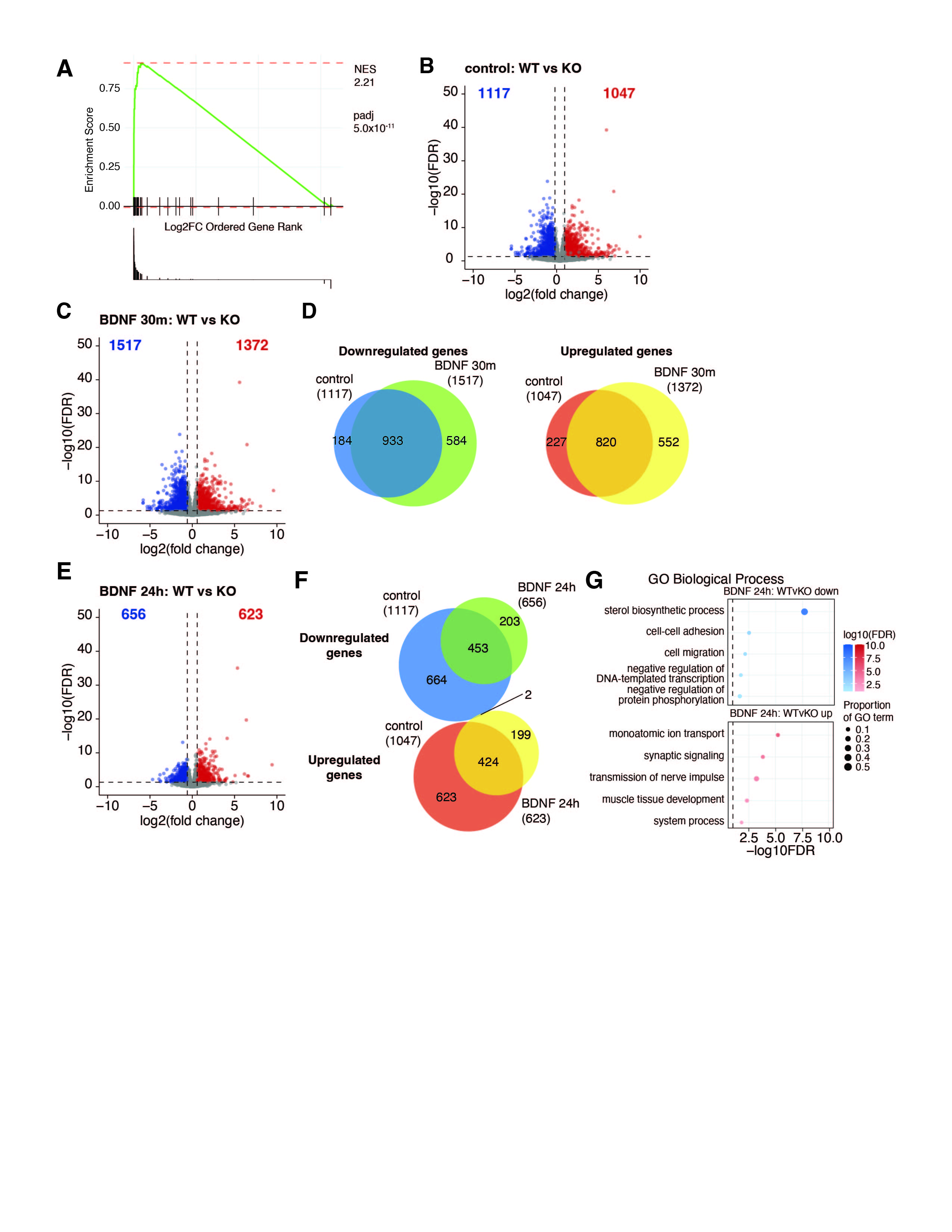

### Supplemental Figure 4

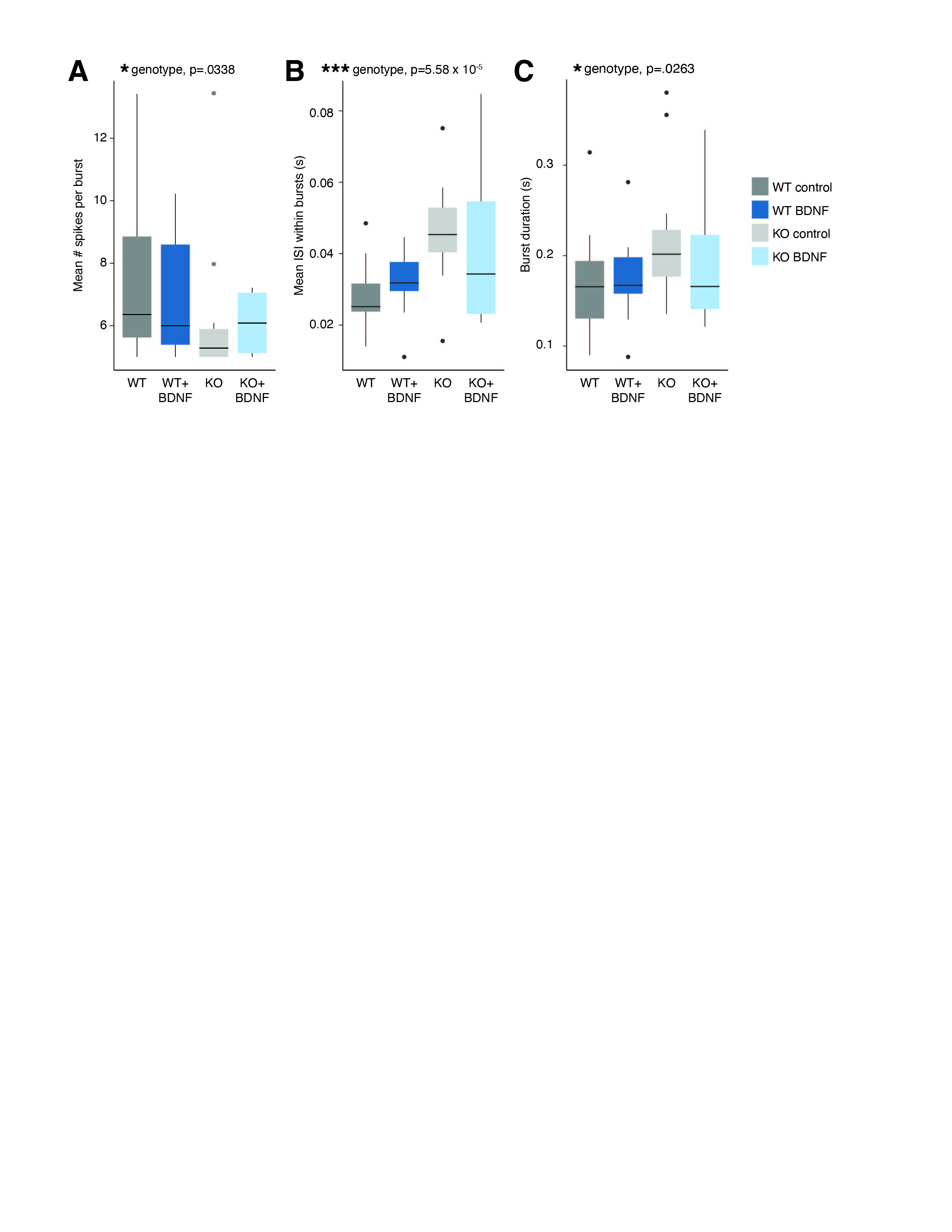

### Supplemental Figure 5

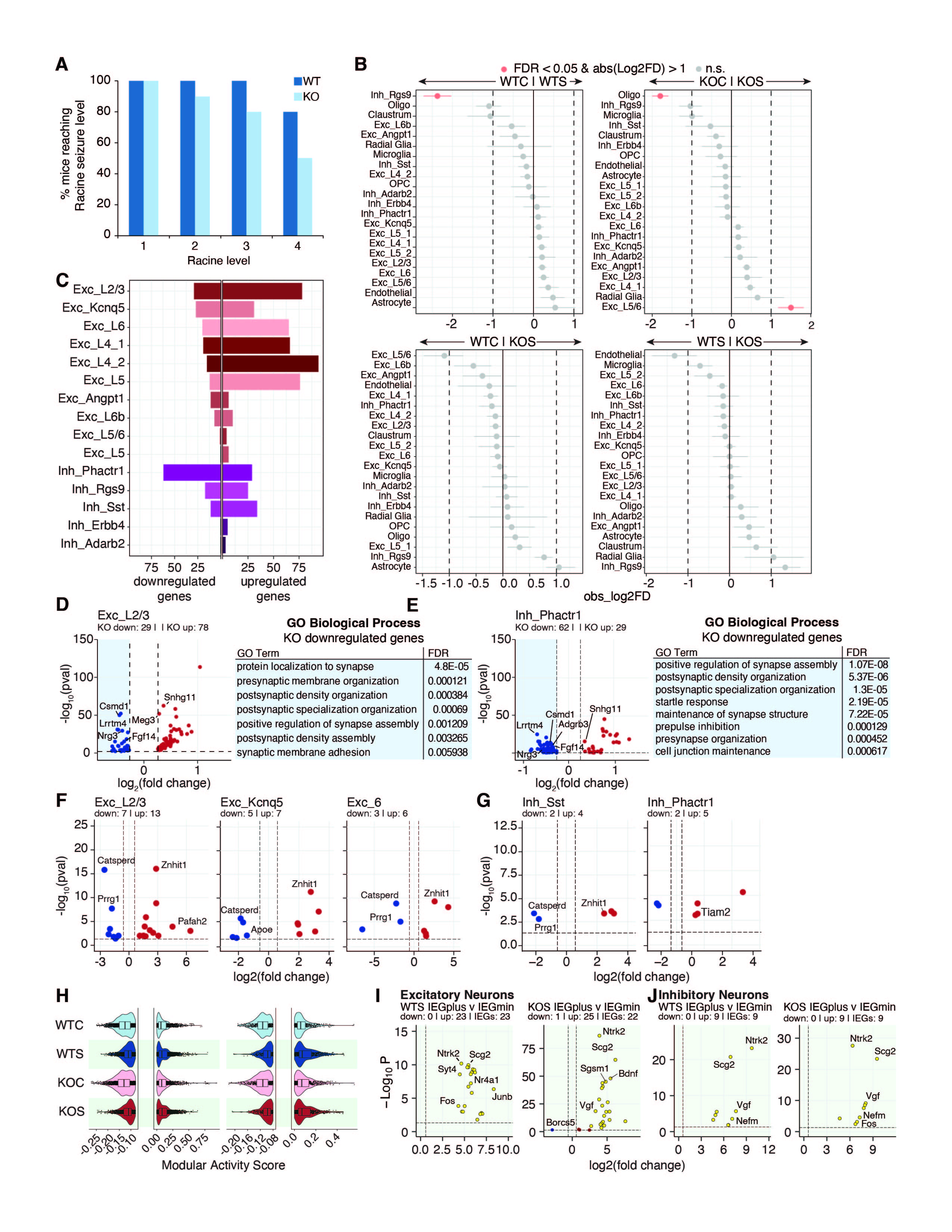
